# Default Mode Network Ventral Hub Connectivity Associated with Memory Impairment in Temporal Lobe Epilepsy Surgery

**DOI:** 10.1101/2020.11.11.377929

**Authors:** Elliot G. Neal, Long Di, You Jeong Park, Austin Finch, Ferdinand Korneli, Stephanie Maciver, Yarema B. Bezchlibnyk, Mike R. Schoenberg, Fernando L. Vale

**Affiliations:** Department of Neurosurgery and Brain Repair, University of South Florida at Tampa, General Hospital, Tampa, FL, USA; Department of Neurology, University of South Florida at Tampa General Hospital, Tampa, FL, USA; Department of Neurosurgery, Medical College of Georgia, Augusta University, Augusta, GA

**Keywords:** epilepsy surgery, epilepsy imaging, source localization, temporal lobe epilepsy, epilepsy prognosis

## Abstract

In patients undergoing surgery for intractable temporal lobe epilepsy, the relationship between the default mode network and patients’ neurocognitive outcome remains unclear. The objective of this study is to employ non-invasive network mapping to identify the relationship between subdivisions of the default mode network and neurocognitive function before and after epilepsy surgery in patients with temporal lobe epilepsy.

Twenty-seven medically patients with medically refractory temporal lobe epilepsy were prospectively enrolled and received resting state functional MRI and neuropsychological testing both pre- and post-operatively. Connectivity within the default mode network was modeled and average connectivity within the networks was calculated.

Higher pre-operative connectivity in the ventral default mode network hub correlated with impaired baseline performance in a visual memory task. Post-operatively, a decrease in ventral but not dorsal default mode network connectivity was correlated with a deterioration of verbal and logical memory after surgery.

Overall, higher connectivity in the ventral default mode network hub was associated with poor memory function in patients with temporal lobe epilepsy both before and after temporal lobe surgery. Pre-operatively, higher ventral connectivity was associated with worse visual function. Post-operatively, decreased connectivity of the ventral and dorsal default mode network was correlated with a greater decrease in logical and verbal memory when compared with the pre-operation baseline. An imbalance in default mode network connectivity towards the ventral stream and more widespread epilepsy networks may be used to predict memory impairments following surgical intervention and may lead to more tailored surgical decision making based on this non-invasive network modeling.

## 1. Introduction

### 1.1 Temporal Lobe Epilepsy

Temporal lobe epilepsy (TLE) is the most common focal epilepsy in adults [1]. Persistence of seizures leads to decline in verbal and visual memory, which may be associated with progressive hippocampal atrophy and it is possible that uncontrolled seizures will lead to deterioration in extratemporal faculties including executive function, attention, psychomotor speed, and general cognitive function [2-9].

### 1.2 Network Analysis in Epilepsy Surgery

Patients with medically refractory TLE may be candidates for potentially curative epilepsy surgery, and decline in memory function seen with persistent seizures can be arrested and possibly reversed with control of seizures following surgical intervention [2]. The role of resting state fMRI (rsfMRI) and network analysis in epilepsy surgery has not been clearly established, but promising data has demonstrated its ability to help lateralize epileptogenesis and predict seizure recurrence after surgery [10-12]. Since uncontrolled seizures in patients with TLE leads to deterioration in extra-temporal neurocognitive function, a more nuanced approach might consider functional connectivity not only within the epilepsy network, but with networks underlying brain function more broadly, such as the default mode network (DMN).

### 1.3 Default Mode Network

The DMN is an intrinsic connectivity network that activates during periods of restful wakefulness – when the brain is not involved in externally oriented tasks – and deactivates during task performance [13, 14]. Functionally, the DMN can be subdivided into two integrated hubs, one more ventral and one more dorsal [15]. In general, studies conducted in patients with TLE have observed decreased connectivity between their temporal lobes and the DMN [16-20]. This finding appears to be related to the duration of TLE [17, 18], and studies correlating data from rsfMRI and tractography suggest that the decreased connectivity may be related to microstructural damage in white matter bundles due to persistent seizures [21]. Combined magnetoencephalography (MEG)/EEG and rsfMRI studies in patients with TLE have shown that during spike-free intervals, connectivity is increased between regions of the temporal lobe and the DMN [22]. Furthermore, left-sided mesial temporal sclerosis (MTS) appears to be associated with decreased functional connectivity of the temporal lobe and the DMN, while increased connectivity is seen in patients with right-sided MTS [22-24]. After surgery, McCormick et al. have shown that the decreased functional connectivity between the contralateral hippocampus and the posterior cingulate cortex (PCC)/precuneus – a critical node within the DMN – predicts postoperative memory decline [25].

### 1.4 Objective

As regions of the DMN are also involved in episodic and autobiographical memory [26, 27], aberrant connectivity within this network may be associated with cognitive impairment in patients with TLE [28]. Here, we more broadly explore DMN connectivity and its correlation with neuropsychological measurements both pre- and post-operatively. The hypothesis for this study is that DMN ventral and dorsal hub connectivity will correlate with memory function both pre- and post-operatively in patients with TLE undergoing surgery.

## 2. Materials and Methods

### 2.1. Patient Demographics

All reported data followed the Strengthening the Reporting of Observational studies in Epidemiology (STROBE) guidelines for observational trials. DMN connectivity and epilepsy networks were modeled in twenty-seven patients with TLE. The patients included in this study represent a consecutive series of patients with TLE who signed consent and agreed to participate in this study (Table 1). The period of data collection started in May 2017 and concluded in October 2020. Each patient underwent a pre-surgical workup for epilepsy surgery including: MRI, long-term video-EEG monitoring (LTM), Wada testing, ^18^Fluoro-2-deoxyglucose positron emission tomography ((^18^F-FDG) PET), and quantitative neuropsychological evaluation. Five of those patients underwent subsequent phase II invasive monitoring for further clarification of epileptogenic focus. EEG and imaging interpretation were performed by a multidisciplinary team blinded to the network modeling parameters. The dominant hemisphere was defined as the hemisphere that supported expressive language when the contralateral side was injected during the Wada test. Additional data points collected included the side of surgery, pathologic MTS diagnosis, and age at surgery.

**Table 1.** Demographics

| Patient Number | Gender | Age at Surgery | MTS (Tissue Specimen) | Surgery Side | Dominant Hemisphere (Wada) | Seizure Free |
| --- | --- | --- | --- | --- | --- | --- |
| 1 | Female | 50 | No | Right | Left | No |
| 2 | Male | 26 | Yes | Left | Left | No |
| 3 | Male | 17 | No | Left | Right | No |
| 4 | Female | 26 | No | Right | Left | No |
| 5 | Female | 35 | No | Left | Left | No |
| 6 | Female | 32 | No | Left | Right | No |
| 7 | Female | 40 | No | Right | Left | No |
| 8 | Female | 36 | No | Left | Left | No |
| 9 | Female | 47 | Yes | Left | Left | Yes |
| 10 | Male | 30 | N/A | Right | N/A | Yes |
| 11 | Male | 23 | N/A | Left | Left | Yes |
| 12 | Female | 34 | Yes | Left | Right | Yes |
| 13 | Female | 58 | No | Left | Left | Yes |
| 14 | Female | 32 | N/A | Left | Bilateral | Yes |
| 15 | Female | 19 | Yes | Right | Left | Yes |
| 16 | Female | 40 | Yes | Right | Left | Yes |
| 17 | Female | 30 | No | Right | Left | Yes |
| 18 | Male | 26 | Yes | Left | Left | Yes |
| 19 | Male | 44 | N/A | Right | Right | Yes |
| 20 | Female | 33 | Yes | Left | Left | Yes |
| 21 | Female | 32 | N/A | Left | Left | Yes |
| 22 | Female | 36 | Yes | Left | Right | Yes |
| 23 | Male | 32 | No | Left | Left | Yes |
| 24 | Female | 28 | Yes | Left | Left | Yes |
| 25 | Female | 24 | No | Left | N/A | Yes |
| 26 | Male | 25 | Yes | Left | Left | Yes |
| 27 | Female | 53 | Yes | Left | Left | Yes |

### 2.2. Data Acquisition

EEG and rsfMRI were obtained on two separate visits. EEG was acquired with twenty-four scalp electrodes in an International 10-20 configuration. rsfMRI was conducted in a three tesla MRI with a blood oxygenation level dependent (BOLD) MRI sequence, consisting of a single five-minute acquisition (eyes closed) with parameters as follows: echo time (TE) of 35 ms, repetition time (TR) of 3000 ms, and a voxel size of 4 x 3.75 x 3.75 mm.

### 2.3. Default Mode Network Connectivity

rsfMRI datasets were normalized to Montreal Neurological Institute (MNI) space using the six-parameter rigid body spatial transformation algorithm using SPM12 (Wellcome Department of Imaging Neuroscience, University College London, UK). An atlas of ROIs generated in a prior study of rsfMRI datasets was overlaid on the rsfMRI to extract the time series signature from regions of interest (ROIs) involved in the ventral and dorsal DMN [13, 29]. An important consideration in this analysis is that of the ROIs used to define the DMN hubs. Whereas other studies have used individual component analyses (ICA) to define the DMN on an individual level [30], in the current study we opted to use an atlas-based, ROI approach. The DMN ROIs were adapted from a previous study using *a priori* methods to identify networks in healthy patients using rsfMRI [29]. The ventral DMN ROI included: a large cluster in the medial parietal cortex, including the precuneus, PCC, and retrosplenial cortex, regions of the bilateral angular gyri, the anterior ventral area of the medial prefrontal cortex as well as bilateral parahippocampal gyri, bilateral inferior temporal cortices, and bilateral superior/middle frontal cortices. Dorsal DMN ROIs included: a major cluster in the medial prefrontal cortex/anterior cingulate cortex (ACC), and the bilateral caudate nuclei. To a lesser degree, the PCC was also included as well as the bilateral hippocampi, thalami, the right angular gyrus, the left superior temporal cortex, and the right calcarine cortex (Figure 1). The average time series for each ROI was used to generate a connectivity matrix of Pearson correlation values grouped into the ventral and dorsal DMN groups. Average connectivity for both the ventral and dorsal DMN was calculated and used as a marker for functional connectivity within the respective network both pre- and post-operatively.

**Figure 1:**
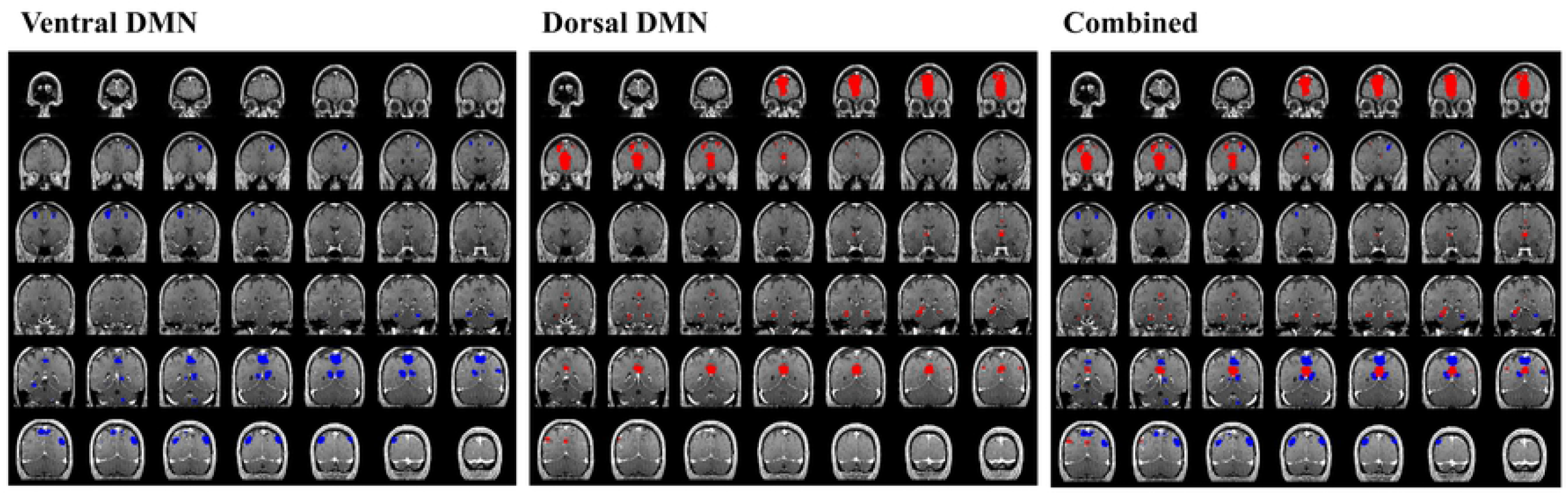
Default Mode Network Regions of Interest: The regions of interest (ROIs) applied for connectivity analysis in the default mode network (DMN) are overlaid on coronal MRI sections. Regions in blue represent the ventral DMN, while those in red represent the dorsal DMN.

### 2.4. Neuropsychological Testing

Pre-operatively, all twenty-seven patients completed a comprehensive neuropsychological assessment following National Institutes of Health (NIH) Epilepsy common data elements recommendations. Due to differences in how one patient’s assessment was documented, the data for that patient could not be included. Thus, twenty-six patients had pre-operative neuropsychological data which were available and were included for analysis. Sixteen of these patients also had post-operative testing. Subtests of the Wechsler

Memory Scale-4th Ed. (WMS-IV) and RAVLT were used to measure verbal immediate memory (LM-I, RAVLT trial 6) and verbal delayed memory (LM-II, RAVLT trial 7) [31]. Visual immediate memory tests included the WMS-IV VR-I subtest and visual delayed memory task including WMS-IV VR-II subtest and the ROCFT-delay task. Letter and semantic verbal fluency tasks including the Controlled Oral Word Association Test (FAS) and animal semantic fluency task was measured. Confrontation naming was measured using the Boston Naming Test (BNT). Executive function including the Wisconsin Card Sorting Test and the Ruff Figural Fluency Test (RFFT). The RFFT provides a measure of nonverbal mental flexibility including unique designs and perseverative errors error ratio. Finally, each patient completed the Wechsler Adult Intelligence Scale – 4th Ed (WAIS-IV) prorated full-scale intelligence index (Table 2). Age-corrected scores for all neuropsychological tests except for WAIS-IV IQ scores were used in analyses. Descriptive statistics for this cohort are given in Table 3.

**Table 2.**
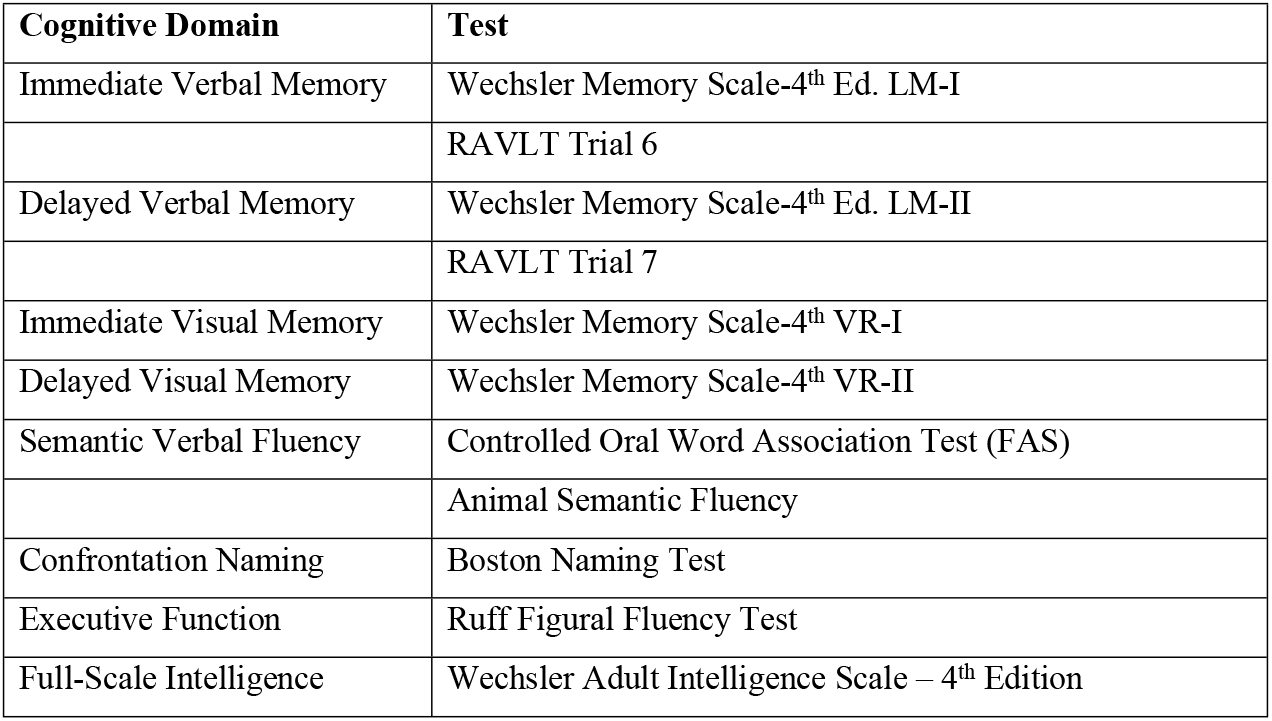
Neuropsychological Tests

| Cognitive Domain | Test |
| --- | --- |
| Immediate Verbal Memory | Wechsler Memory Scale-4<sup>th</sup> Ed. LM-I |
|  | RAVLT Trial 6 |
| Delayed Verbal Memory | Wechsler Memory Scale-4<sup>th</sup> Ed. LM-II |
|  | RAVLT Trial 7 |
| Immediate Visual Memory | Wechsler Memory Scale-4<sup>th</sup> VR-I |
| Delayed Visual Memory | Wechsler Memory Scale-4<sup>th</sup> VR-II |
| Semantic Verbal Fluency | Controlled Oral Word Association Test (FAS) |
|  | Animal Semantic Fluency |
| Confrontation Naming | Boston Naming Test |
| Executive Function | Ruff Figural Fluency Test |
| Full-Scale Intelligence | Wechsler Adult Intelligence Scale – 4<sup>th</sup> Edition |

**Table 3.** Neuropsychological Testing Values

| **Test Name** |  | **Minimum** | **Maximum** | **Mean** | **Standard Deviation** |
| --- | --- | --- | --- | --- | --- |
| **FSIQ** | Pre-op score | 64.00 | 112.00 | 85.50 | 14.70 |
|  | Post-op score | 61.00 | 118.00 | 84.86 | 15.27 |
|  | Difference score | -18.00 | 16.00 | 1.08 | 7.99 |
| **LM-I** | Pre-op score | 20.00 | 64.00 | 37.83 | 13.66 |
|  | Post-op score | 20.00 | 59.00 | 36.79 | 13.47 |
|  | Difference score | -29.00 | 10.00 | -3.46 | 10.65 |
| **LM-II** | Pre-op score | 20.00 | 66.00 | 38.78 | 12.26 |
|  | Post-op score | 20.00 | 59.00 | 37.80 | 11.77 |
|  | Difference score | -15.00 | 13.00 | -1.92 | 7.69 |
| **VR-I** | Pre-op score | 20.00 | 63.00 | 38.70 | 12.70 |
|  | Post-op score | 19.00 | 60.00 | 36.87 | 12.42 |
|  | Difference score | -23.00 | 17.00 | -0.69 | 10.23 |
| **VR-II** | Pre-op score | 20.00 | 73.00 | 40.70 | 11.17 |
|  | Post-op score | 20.00 | 66.00 | 39.13 | 14.81 |
|  | Difference score | -12.00 | 22.00 | 1.85 | 9.33 |
| **FAS** | Pre-op score | 21.00 | 60.00 | 37.56 | 10.71 |
|  | Post-op score | 23.00 | 58.00 | 39.33 | 7.90 |
|  | Difference score | -7.00 | 19.00 | 4.57 | 6.91 |
| **Animal Naming** | Pre-op score | 24.00 | 60.00 | 39.20 | 9.60 |
|  | Post-op score | 17.00 | 53.00 | 33.93 | 10.61 |
|  | Difference score | -26.00 | 12.00 | -2.79 | 11.05 |
| **RAVLT-6** | Pre-op score | 0.00 | 80.00 | 40.82 | 17.23 |
|  | Post-op score | 0.00 | 57.00 | 36.75 | 16.88 |
|  | Difference score | -22.00 | 21.50 | 0.50 | 11.58 |
| **RAVLT-7** | Pre-op score | 5.00 | 64.00 | 38.24 | 15.19 |
|  | Post-op score | 10.00 | 56.00 | 37.71 | 16.05 |
|  | Difference score | -14.00 | 31.00 | 4.46 | 12.76 |
| **BNT** | Pre-op score | 8.00 | 53.00 | 34.92 | 11.08 |
|  | Post-op score | 8.00 | 53.00 | 33.87 | 11.49 |
|  | Difference score | -15.00 | 15.00 | -1.14 | 8.29 |
| **RFFT-UD** | Pre-op score | 33.00 | 120.00 | 67.76 | 22.40 |
|  | Post-op score | 42.00 | 116.00 | 68.27 | 20.23 |
|  | Difference score | -17.70 | 12.00 | 0.61 | 7.51 |
| **RFFT-ER** | Pre-op score | 36.00 | 73.00 | 55.51 | 14.27 |
|  | Post-op score | 38.00 | 73.00 | 55.83 | 10.93 |
|  | Difference score | -21.00 | 9.20 | -3.47 | 8.91 |

### 2.5. Statistical Analysis

Neuropsychological data and the network metrics were compared using a Spearman Rho correlation coefficient analysis. Connectivity differences between the subgroups was analyzed using an independent-sample t-test. All statistical tests were conducted using IBM SPSS Statistics Version 26 (IBM Corp., Armonk, New York, United States). P-values less than α = 0.05 were considered significant.

### 2.6. Data Availability

The data will be made available to anyone within reason who requests it from the corresponding author. The network mapping algorithm will also be made available for purposes of validation and corroboration of presented results if requested.

## 3. Results

### 3.1. Demographics

Twenty-seven patients with TLE underwent pre-operative rsfMRI scanning and network analysis. The time from the first lifetime seizure to the surgery was an average of 14.9 +/- 10.2 years. The average time to most recent follow-up after surgery was 30.2 +/- 8.69 months. Nineteen (70%) of the patients underwent surgery on the left temporal lobe. Twenty-five patients (93%) completed Wada testing, and fifteen of those patients (60%) had a surgery in the dominant hemisphere. In the cohort, four patients underwent stereotactic laser amygdalohippocampotomy (SLAH) while the remaining twenty-three underwent microsurgical resection with either selective amygdalohippocampectomy (SAH; sixteen), anterior temporal lobectomy (ATL; one), or resection of the temporal pole with amygdalectomy and minimal hippocampal resection (HC-sparing; six). Of these patients, eleven (41%) had tissue specimen proven MTS, eleven (41%) did not have MTS, and five (19%) patients had no hippocampus specimens collected.

### 3.2. Pre-operative DMN Connectivity in TLE

Pre-operative connectivity within the ventral and dorsal DMN and the ratio of ventral to dorsal DMN connectivity were compared to pre-operative neuropsychometric performance. To control for possible differences in DMN connectivity between patients, we analyzed the relationship between the ratio of ventral to dorsal DMN connectivity. Patients with a higher ventral:dorsal DMN connectivity ratio (DCR; i.e. those with a relatively higher ventral DMN connectivity when compared to dorsal DMN connectivity) performed worse in immediate (VRI I, R = −0.401, p = 0.042) and delayed (VRI II, R = −0.446, p = 0.022) visuoconstructional memory tasks. Higher DCR (ventral > dorsal) also correlated with impaired immediate and delayed verbal memory function (RAVLT6, R = −0.434, p = 0.024; RAVLT7 R = −0.383, p = 0.049) (Figure 2).

**Figure 2:**
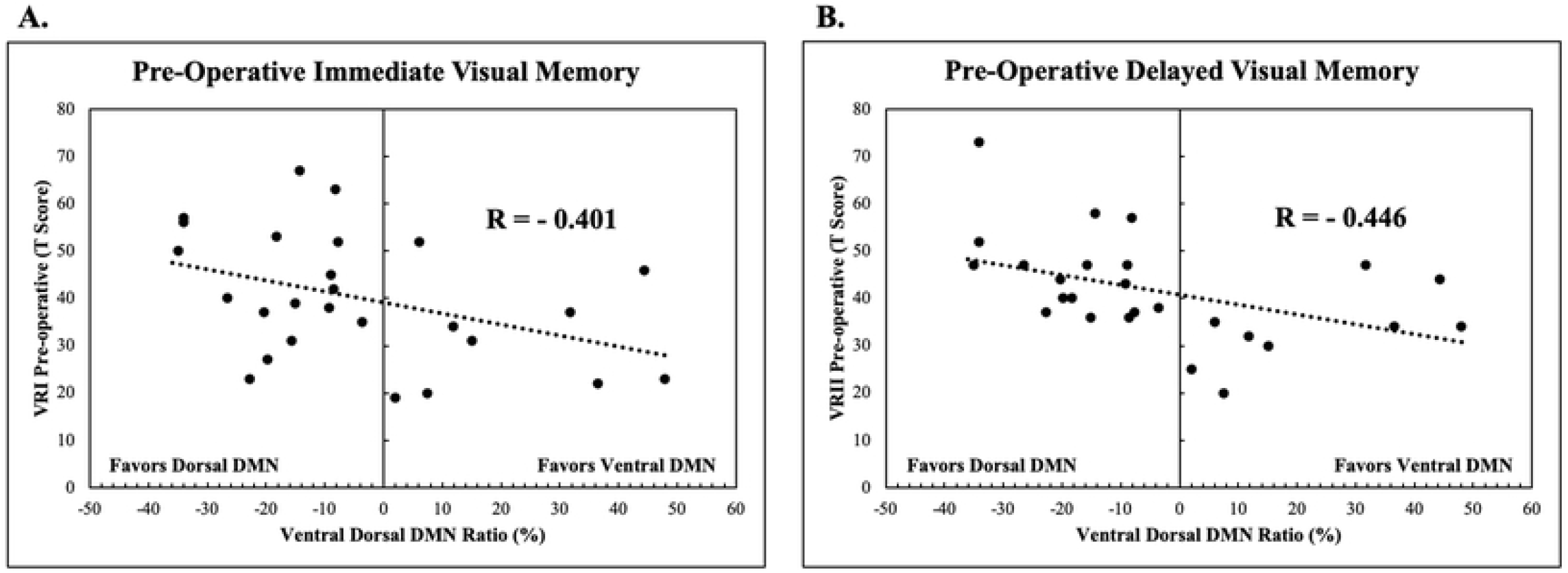
Increased pre-operative DMN ventral hub connectivity correlates with relatively worse visual memory. A. Immediate visual memory (LMI) pre-operative performance, scaled to age-matched controls, is lower in patients who’s DMN connectivity is relatively higher in the ventral hub compared to the dorsal hub. B. The same trend is also found when comparing the ratio of ventral to dorsal DMN hub connectivity to delayed visual memory. The connectivity ratio is skewed towards the ventral hub in patients who performed worse on the delayed visual memory task (LMII).

### 3.3. Post-operative Changes in the DMN

DMN connectivity and neuropsychological function were also measured post-operatively in the same way as the pre-operative assessment. Change in connectivity of both the ventral and dorsal DMN after surgery was measured by the ratio of the average pre- and post-operative Pearson correlation within the respective network. For example, when the post-operative network connectivity was close to that of the pre-operative network (ratio approaches unity), then that patient’s network connectivity was relatively preserved after surgery. Pre-operative ventral DMN connectivity was compared to post-operative ventral DMN connectivity, and pre-operative dorsal DMN was compared to post-operative dorsal DMN. It was found increase in connectivity within the ventral DMN after surgery was associated with a decline post-operatively in both immediate (LM I, R = −0.668, p = 0.006) and delayed (LM II, R = −0.747, p = 0.001) (RAVLT 7, R = −0.622, p = 0.013) verbal memory (Figure 3).

**Figure 3:**
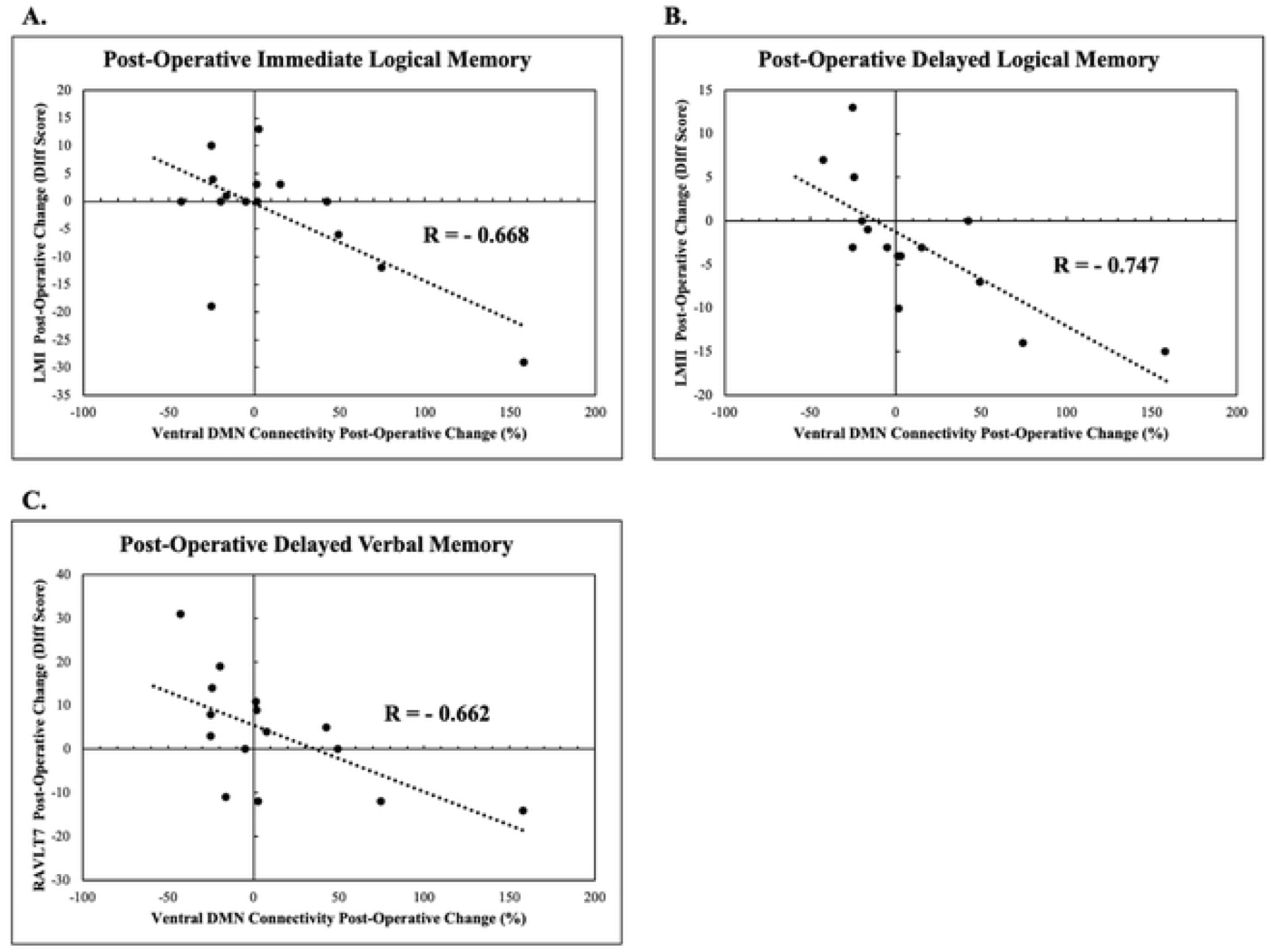
Post-operative connectivity in the ventral hub is increased in patients who exhibit a relative decline in memory compared to their pre-operative performance. Difference scores are calculated by subtracting the pre-operative score from the post-operative score so that higher difference scores indicate improvement post-operatively compared to before surgery. Increased connectivity within the ventral DMN hub correlated with decline in immediate logical memory (A), delayed logical memory (B), and delayed verbal memory (C).

### 3.4. Subgroup Analysis

DMN connectivity was also assessed in patients segregated by several factors, including presence of MTS, left vs. right side surgery, and dominant vs. non-dominant side surgery. First, comparison was made with regards to the pre-operative DMN connectivity. When comparing patients with pathology-proven MTS to those without (n=x, y, respectively), neither ventral nor dorsal DMN connectivity at baseline were significantly different (ventral p = 0.754, dorsal p = 0.815). The same results were found when comparing patients with surgery on the left side vs. the right (n=x, y, respectively; ventral p = 0.722, dorsal p = 0.366) and between patients with surgery on the dominant vs. non-dominant side (n=x, y, respectively; ventral p = 0.626, dorsal p = 0.738). At the post-operative timepoint, again there was no difference in ventral or dorsal DMN connectivity with respect to presence of MTS on pathology (ventral p = 0.603, dorsal p = 0.282), side of surgery (ventral p = 0.182, dorsal p = 0.690), or surgery on the dominant vs. non-dominant side (ventral p = 0.544, dorsal p = 0.721). Pathological specimens were not available for five patients, so the analysis was also performed based on radiographic features of MTS. Again, there were no significant differences in the ventral or dorsal DMN connectivity either pre-operatively (n=x, y, respectively; ventral DMN p = 0.646, dorsal DMN p = 0.938) or post-operatively (n=x, y, respectively; ventral DMN p = 0.684, dorsal DMN p = 0.323).

## 4. Discussion

### 4.1. Main Findings and Impact

In the present study, we show that increased ventral hub connectivity in patients with medically refractory temporal lobe epilepsy was correlated with impaired memory function both before and after temporal lobe surgery. Pre-operatively, we found that increased ventral, but not dorsal, DMN connectivity in patients with TLE is associated with poorer immediate and delayed verbal and visual. After surgery, it was shown that relative increase in connectivity within the ventral DMN was associated with a decrease in immediate and delayed logical memory and delayed verbal memory. Subgroup analyses revealed no difference in DMN connectivity either pre-operatively or post-operatively in patients with pathological or radiological diagnoses of MTS, patients undergoing surgery on the left or right side, or patients undergoing dominant or non-dominant hemisphere surgeries.

Here, increased connectivity within the ventral DMN at baseline was generally found to be a poor prognostic indicator in that it was associated with impaired verbal and visual memory pre-operatively and relative increase in ventral DMN connectivity after surgery was associated with a greater decline in verbal and logical memory. These findings may relate to the more global effects on network connectivity seen in patients with TLE. One possible explanation is that with longstanding epilepsy, the epileptogenic temporal lobe becomes progressively disconnected from the DMN over time [16-20], resulting in an associated dysregulation in the DMN which is measured as an increase in functional connectivity of the ventral stream of the DMN, perhaps by way of compensation. The dysregulated ventral DMN may then become less “resilient” to surgical interventions and may result in it becoming abnormally hyperactivated after surgery, with an end effect in impairments of memory and visuoconstructional abilities.

Interestingly, we failed to identify any significant differences between patients with either pathological or radiographic diagnoses of MTS vs. no-MTS, left vs. right sided surgery, or dominant vs. non-dominant surgery side. The lack of an association is not clear, but an unknown compensatory network may play an important role in these patients. One prior study done in patients with MTS found a decreased connectivity between the PCC, precuneus, and mesial temporal lobes, but there was no comparison of patients with MTS and those without MTS [21].

In future studies, we will include stereo-encephalography (SEEG) results into this analysis to incorporate seizure spread as it relates to the DMN hubs. We do not yet know how seizure propagation disrupts the DMN, and different patterns of spread may help to explain why some patients have a more dysregulated ventral hub than others. Furthermore, semiology has been related to anatomy [32, 33], but not yet to network connectivity analysis. In future studies we will incorporate semiology into this analysis to differentiate subgroups of patients who may more preferentially have disrupted DMN hub connectivity.

While not detracting from the findings of the present study, there are some limitations. First, the sample size is somewhat limited. This is a preliminary study showing that is intended to inform larger prospective studies into the DMN connectivity in patients undergoing temporal lobe surgery for epilepsy. Furthermore, while the sample size is limited, follow up is rigorous including post-operative neuropsychological testing and rsfMRI. Another limitation of this study is that only one method is used to assess connectivity, and in future studies we hope to use SEEG to corroborate these connectivity findings.

Connectivity within the DMN was investigated in patients with TLE undergoing surgery both pre- and post-operatively using rsfMRI. Increased connectivity within the ventral DMN was found to be a poor prognostic indicator in that it was associated with worse visual and verbal memory pre-operatively, and increased connectivity in the ventral DMN after surgery was found in patients who had relatively more decline in verbal and logical memory post-operatively.

## 5. Acknowledgements

None

## Abbreviations

((^18^F-FDG) PET): ^18^Fluoro-2-Deoxyglucose Positron Emission Tomography
(BOLD): Blood Oxygenation Level Dependent
(BNT): Boston Naming Test
(COWAT-FAS): Controlled Oral Word Association Test
(DMN): Default Mode Network
(EEG): Electroencephalography
(LTM): Long-Term Video-EEG Monitoring
(MEG): Magnetoencephalography
(MTS): Mesial temporal sclerosis
(MNI): Montreal Neurological Institute
(NIH): National Institutes of Health
(rsfMRI): Resting state functional MRI
(RAVLT6): Rey Auditory Verbal Learning Test, Trial 6
(RAVLT7): Rey Auditory Verbal Learning Test, Trial 7
(RFFT-UD): Ruff Figural Fluency Test - Unique Designs
(RFFT-ER): Ruff Figural Fluency Test – Error Ratio
(SPECT): Single-Photon Emission Computed Tomography
(TLE): Temporal Lobe Epilepsy
(FSIQ): Wechsler Adult Intelligence Scale-4^th^ Ed. – Full Scale Intelligence Quotient
(WMS-IV): Wechsler Memory Scale-4^th^ Ed.
(LM-I): Wechsler Memory Scale-4^th^ Ed., Logical Memory Immediate recall subtest
(LM-II): Wechsler Memory Scale-4^th^ Ed., Logical Memory Delayed recall subtest
(VR-I): Wechsler Memory Scale-4^th^ Ed., Visual Reproduction Immediate Recall subtest
(VR-II): Wechsler Memory Scale-4^th^ Ed., Visual Reproduction Delayed Recall subtest

